# Simultaneous proteomics and three PTMomics characterization of pro-inflammatory cytokines stimulated INS-1E cells using TiO_2_ enrichment strategy

**DOI:** 10.1101/509125

**Authors:** Honggang Huang, Lylia Drici, Pernille S. Lassen, Giuseppe Palmisano, Martin R. Larsen

**Affiliations:** Department of Biochemistry and Molecular Biology, University of Southern, Denmark; The Danish Diabetes Academy, Odense, Denmark; Arla Foods Ingredients Group P/S, Denmark; Department of Parasitology, ICB, University of São Paulo, Brazil

**Keywords:** PTMomics, Phosphorylation, Reversible Cys modifications, Sialylated N-linked glycosylation, TiO_2_ enrichment, Cytokines, INS-1E cells, Diabetes

## Abstract

Diverse protein post-translational modifications (PTMs) in proteins form complex combinatorial patterns to regulate the protein function and biological processes in a fine-tuning manner. Reversible phosphorylation, cysteines (Cys) modification, and N-linked glycosylation are essentially involved in cellular signaling pathways of pro-inflammatory cytokines, which can induce beta cell death and diabetes. Here we developed a novel mass spectrometry–based proteomic strategy (termed TiCPG) for the simultaneous comprehensive characterization of the proteome and three post-translational modifications (PTMomes) by applying TiO_2_ enrichment of peptides with reversibly modified Cysteine (rmCys), Phosphorylation, and sialylated N-linked (SAN-) Glycosylation from low amount of sample material with largely minimized sample loss. We applied this TiCPG strategy to quantitatively study the change of the three PTMs in β-cell-like INS-1E cells subject to pro-inflammatory cytokines stimulation. It enabled efficient enrichment and quantitative analysis of 8346 rmCys sites, 10321 phosphosites and 1906 SAN-glycosylation sites from 5853 proteins. Significant regulation was found on 100 proteins at the total protein level, while much higher degree of regulation was identified on 3025 peptides with PTMs from 1490 proteins. The three PTMs were co-regulated in proteins, but demonstrated differential spatial and temporal patterns related to protein cellular localization and function in the time course of cytokines stimulation, and they were extensively involved in essential signaling pathways related to pro-inflammatory cytokine mediated β-cell apoptosis, such as the inducible NO synthase (NOS2) signaling pathway, Overall, the TiCPG strategy is proved as a straight forward and powerful tool for multiple PTMomics studies.

## Introduction

Protein post-translational modifications (PTMs) represent an essential regulatory mechanism for protein function. They act as molecular switches through the modulation of numerous properties of proteins including enzymatic activity, protein interactions and subcellular location. Cells in response to internal and external changes, such as stimulation and stress, can be rapidly relayed from sensors to effectors via reversible PTMs of proteins. There are more than 450 PTMs listed in the Uniprot database [1], Diverse PTMs must be fine-tuned in the regulatory network in order to coordinate the protein states, and synergistically regulate the protein function in specific cellular conditions in a complex and dynamic way [2, 3]. Phosphorylation, cysteine (Cys) modification, and Sialylated N-linked (SAN-) glycosylation are reversible PTMs essentially involved in cellular signaling networks and pathways. These three PTMs are highly represented in the proteome, they often co-exist on many proteins, and there is a possibility for cross-talking between these PTMs in order to regulate and fine-tune cellular signaling. For example, the oxidation of the catalytic Cys residues in the HC(X)5R motif of protein tyrosine phosphatases (PTPs) can completely abolish PTPs activity for tyrosine dephosphorylation [4], indicating the crosstalk between phosphorylation and Cys oxidation. The modulation of phosphorylation events through cell membrane sialylation has been described in cancer [5, 6], where tumors from sialyltransferase-deficient animals displayed decreased phosphorylation of focal adhesion kinase, which was involved in the signaling pathway of β1-integrins[7].

Infiltration of the islet of Langerhans by immune cells is a common phenomenon observed in type 1 and 2 diabetes. Elevated numbers of immune cells directly cause increased levels of inflammatory cytokines and chemokines in islets of diabetic patients and animal models for diabetes [8-10]. Tumor necrosis factor-α (TNF-α), interferon-γ (IFN-γ) and interleukin 1β (IL-1β) are among the primary components of pro-inflammatory cytokines responsible for the inflammatory response in diabetes pathogenesis [11]. They can increase the production of reactive oxygen species (ROS) through mitochondria and NADPH oxidase [12, 13]. The production of nitric oxide (NO) was largely increased by the stimulation of cells with IFN-γ/TNF-α through the increased expression of inducible NO synthase (iNOS or NOS2) [14]. The ROS can induce apoptosis and impair beta-cell function and viability in type 1 and 2 diabetes [15, 16]. Cys is the most ROS-sensitive amino acid, which undergoes a variety of reversible regulatory oxidative PTMs mediated by ROS [17], and as such, Cys oxidation should be greatly affected in beta-cells subjected to cytokines stimulation. However, ROS can also affect other PTMs such as phosphorylation and glycosylation. ROS can induce increased serine phosphorylation but decreased tyrosine phosphorylation of insulin receptor substrate protein 1 (IRS1) and other key proteins, which results in enhanced protein degradation and impaired insulin signaling [12]. Interestingly, in yeast, defects in N-glycosylation lead to production of ROS and apoptosis [18]. Studies have revealed that IFN-γ/TNF-α co-stimulation induced beta-cell apoptosis, partially through the regulation of protein expression and phosphorylation involving the Janus kinase - signal transducer and activator of transcription (JAK-STAT) and the nuclear factor-kappa B (NF-κB) pathways, and through increased nitric oxide production in human pancreatic islets[19-21]. However, the underlying molecular mechanisms of cytokines mediated beta-cell apoptosis, including site specific PTM modulation, are largely unknown.

Advanced mass spectrometry (MS) in combination with various PTM-specific enrichment methods enable the high-throughput identification and characterization of PTMs in cells or tissues. So far, most large-scale studies have focused on analyzing a single PTM; however, the analysis of the interaction between multiple PTMs requires the use of multiple complementary or comprehensive enrichment strategies for multiple PTMs[2, 3]. The Titanium Dioxide (TiO_2_) beads have a high affinity capacity for adsorption of the acidic groups, such as the phosphate-group, to the surface of TiO_2_ by forming a bridging bidentate binding [22]. Based on this principle, we have previously developed several enrichment methods for different PTMs using TiO_2_ resin. The method for enrichment of phosphopeptides using TiO_2_ is widely used for phosphoproteomic studies with high efficiency and specificity [23]. Meanwhile, the TiO_2_ method was also developed for enrichment of SAN-glycopeptides [24]. Recently, we developed a new TiO_2_-based method for enrichment of cysteine-containing peptides (Cys peptides) selectively labelled with a novel synthesized Cysteine-specific Phosphonate Adaptable Tag (CysPAT) [25, 26]. Taken together, these methods indicate the possibility of developing a comprehensive strategy using TiO_2_ for simultaneous enrichment of phosphopeptides, reversibly oxidized Cys peptides and sialylated N-linked glycopeptides and the simultaneous characterization of these three PTMs in biological settings.

The pro-inflammatory cytokines TNF-α and IFN-γ can increase the production of reactive oxygen species (ROS) and nitric oxide (NO) in β-cells through the increased expression of inducible NO synthase (iNOS or NOS2) and induce apoptosis or impair β-cell function and viability in type 1 and 2 diabetes [14]. The oxidative status of Cys, phosphorylation and glycosylation could be significantly regulated in this process, but it is largely unknown on a global scale. Therefore, in this study, we combined the three TiO_2_ enrichment methods to develop a comprehensive TiCPG strategy for simultaneous analysis of the proteome and the three abovementioned PTMomes from low amount of starting material, and applied this multiple PTMomics strategy to quantitatively study the change of proteome and these three PTMomes, and their interaction in insulinoma pancreatic beta-cell line, INS-1E cells subject to IFN-γ/TNF-α co-stimulation using iTRAQ and LC-MS/MS based quantitative proteomics.

## Results and Discussion

### The TiO_2_-based simultaneous enrichment of reversibly modified Cys peptides, Phosphopeptides, and sialylated N-linked Glycopeptides (TiCPG) strategy

Diverse PTMs form complex combinatorial patterns and cooperate to specify downstream biological processes in a fine-tuning manner. The combined characterization of different PTMs within the same sample is highly desired for systemically analyzing PTMs in biological contexts and deciphering multiple PTM codes [3, 27]. In the past decade, several different TiO_2_ based methods were developed by our group for enrichment of phosphopeptides [23, 28], Sialylated N-linked glycopeptides [24, 29], and Cys peptides [25] with high specificity and efficiency. Based on these methods, we here developed a comprehensive strategy for characterization of the above mentioned three PTMs, the TiO_2_-based simultaneous enrichment of rmCys peptides, Phosphopeptides, and sialylated N-linked Glycopeptides (TiCPG strategy), which for the first time offers the possibility for combined analysis of three different PTMs from a low amount of biological sample. The TiCPG strategy was applied to study the change of the three PTMs in the insulinoma cell line INS-1E cells subjected to 0h, 12h and 24h IFN-γ/TNF-α co-stimulation using iTRAQ based quantitation (Figure 1). Briefly, the INS-1E cell control and the cells subject to 12 h and 24 h IFN-γ/TNF-α co-stimulation were lysed by sonication in cell lysis buffer containing NEM to block the free Cys. The extracted proteins were purified with 10KDa filters to remove the extra NEM, After reduction, tryptic digestion and iTRAQ labeling with100 µg of peptides from each condition (0 hour/control, iTRAQ-114; 12 hours, iTRAQ-115; 24 hours, iTRAQ-116). The mixed 300 µg peptides were treated with the CysPAT to label the reduced Cys on peptides. Subsequently, the peptide mixtures were subjected to TiO_2_ enrichment. The phosphopeptides, CysPAT labeled rmCys peptides and sialylated N-linked glycopeptides were co-enriched in this initial step, the TiO_2_ unbound peptides were collected as they contained the non-modified peptides and represented the general proteome. The co-enriched peptide fraction was further subject to deglycosylation with PNGase F and Sialidase A, resulting in the enzymatic conversion of Asn to Asp (mass increase of 0.9840 Da) at the former site of N-linked glycan attachment with the consensus motif N-X-S/T/C (X ≠ P). After deglycosylation, the co-enriched peptides with the three PTMs can be directly subject to LC-MS/MS analysis with good identification efficiency (data unpublished). However, consider the low abundance of sialylated glycopeptides compared to phosphopeptides and CysPAT labeled Cys peptides in a sample not enriched for membrane proteins, a second round of TiO_2_ enrichment was performed to separate the formerly sialylated N-linked glycopeptides from the co-enriched fraction. Additionally, this step can also further remove potential peptides non-specific binding to TiO_2_ and increase the enrichment specificity for phosphopeptides and CysPAT labeled Cys peptides. After the second round of TiO_2_ enrichment, we obtained three peptide fractions, the non-modified peptides fraction (NM fraction), the formerly sialylated N-linked glycopeptides fraction (G fraction), and the CysPAT labeled Cys peptides and phosphopeptides fraction (CP fraction). All these fractions were pre-fractionated with HILIC, as HILIC was confirmed to be a powerful tool for sub-separation and decrease the complexity of peptide mixture, especially for the fraction containing both the phosphopeptides and CysPAT labeled Cys peptides [25], as phosphopeptides are more hydrophilic than CysPAT labeled Cys peptides. Finally, all the samples were analyzed by LC-MS/MS using the Orbitrap Fusion Tribrid™ mass spectrometer. The experiment was performed in triplicates.

**Figure 1:**
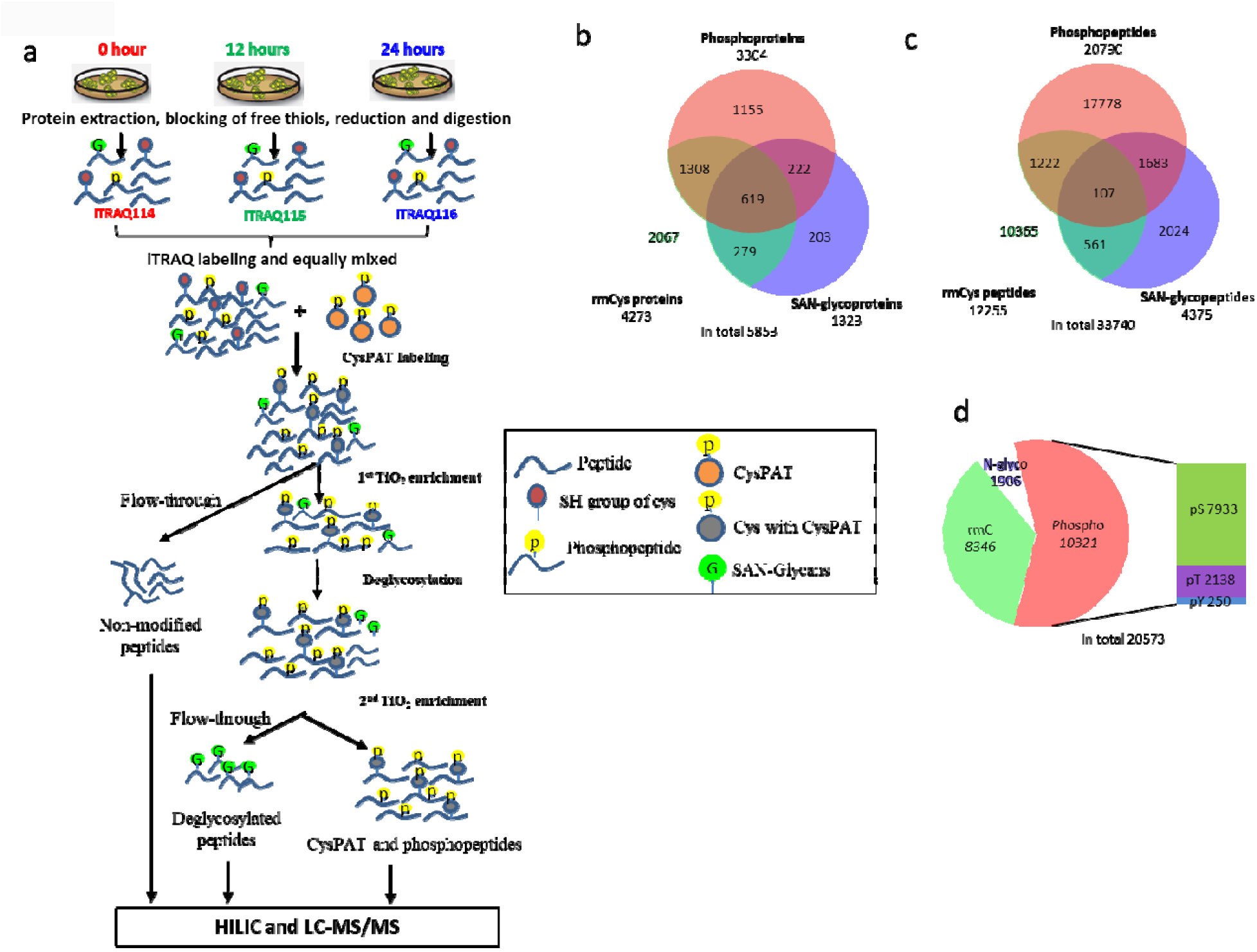
The application of TiCPG strategy for quantitative characterization of INS-1E cells subject to cytokines stimulation enables simultaneous enrichment of rmCys peptides, phosphopeptides and SAN-glycopeptides with high efficiency. (a) The experimental workflow of the TiCPG strategy. (b) The Venn diagram of identified rmCys proteins, phosphoproteins and SA N-glycoproteins from enriched fractions. (c) The Venn diagram of identified unique peptides carrying rmCys, phosphorylation and SAN-glycosylation. (d) The distribution of identified PTM sites.

Currently, three strategies are available for combined analysis of multiple PTMs, the parallel enrichment of PTMs, the serial enrichment of multiple PTMs and top-down analysis of PTMs on the intact protein [3]. These strategies all have some disadvantages and limitations. In principle, the parallel and serial enrichment strategies employ different PTMs enrichment methods for the same sample, and do the individual PTM enrichment separately, therefore, the complete experiment procedures are very complicated and time-consuming, meanwhile, they can not identify peptides with different PTMs. Specifically, the parallel enrichment requires a large amount of starting material for each enrichment experiments, and the serial enrichment of PTMs can be prone to a more pronounced sample loss during the experimental procedure. For top-down analysis of PTMs within intact protein, the sensitivity and techniques are still a great challenging. A strategy for simultaneous enrichment of multiple PTMs may partially overcome these challenges. The TiCPG strategy can offer such solutions with the following advantages. Firstly, the strategy can be used for simultaneous enrichment of phosphopeptides, rmCys peptides and sialylated N-linked glycopeptides in just one TiO_2_ enrichment step, the experimental procedure is straightforward, easy and fast to perform and minimize the problem of sample loss. Secondly, the strategy can detect different forms of peptides unbiasedly, no matter peptides with single PTM or the peptides that contain multiple and/or different types of PTMs on the same peptide, this feature is essential for characterization of PTM cross-talking. In addition, the strategy does not require large amount of starting material, the needed material can be the same as that for single PTM study. In this study, the relatively low amount of starting material (100 µg from each condition, 300 µg in total) with multiplexed iTRAQ isotopic labeling strategy makes it an efficient and attractive approach for PTM studies with limited sample amount, such as the valuable clinical samples. Lastly, the cost of this strategy is relatively low, compare to commercial antibody and resin based methods. Regarding the complexity of co-enriched PTM peptides, the second TiO_2_ enrichment to separate the deglycosylated peptides and the followed HILIC pre-fractionation can greatly decrease the complexity of all samples. The application of advanced high resolution accurate MS is also helpful for increasing the number of peptide identifications. In summary, the TiCPG strategy is an efficient and easy tool for simultaneous characterization of reversible Cys modification, phosphorylation, and N-linked Glycosylation from low amount of biological samples.

### Overall identification and motif analysis

From the NM fractions, we identified 36031 unique peptides derived from 6396 proteins (Supplemental Table S1), and from the enriched fractions, a total of 46593 unique peptides from 6476 proteins were identified (Supplemental Table S2). Among the enriched peptides, 33740 peptides from 5853 proteins contained PTM sites. The Venn diagrams (Figure 1b and Figure 1c) indicated the overlap of proteins and peptides identified with the three PTMs. For these 33740 unique peptides with PTM sites, 3573 peptides were identified with at least 2 different PTMs, and 107 peptides with all three PTMs (Supplemental Table S3, Figure 1c). About 10.6% of the modified peptides identified by the TiCPG strategy contained two or three PTMs, which is much higher than what identified by serial enrichment (0.3%)[30]. About 90% of the enriched peptides were phosphopeptides and rmCys peptides in the CP fractions, indicating high enrichment efficiency, and more than 61.6% of the unique peptides could be identified in at least two biological replicates, indicating a relatively high reproducibility (Supplemental Figure 1). Through retrieving the Uniprot database, 20573 PTM sites were successfully mapped to proteins, including 10321 phosphosites (7933 pS, 2138 pT and 250 pY), 8346 rmCys sites and 1906 formerly SAN-glycosites (Supplemental Table S4, Figure 1d). These results confirmed that the phosphopeptides, the CysPAT labeled rmCys peptides and the formerly SAN-glycopeptides can be simultaneously enriched and characterized by the TiCPG method with high efficiency and sensitivity.

Motif analysis of PTM sites using Motif-X (P < 10^−6^ and relative occurrence rate threshold of 3%) revealed 14 rmCys motifs, 12 of them were formed by rmCys sites with an adjacent lysine (K) or arginine (R) amino acids. It also revealed 12 phosphoserine motifs, 10 phosphothreonine motifs, 1 phosphotyrosine motif and the conserved N-linked glycosylation NXS/T/C motifs. The identified motifs and statistical information were listed in Supplemental table S5.

### Quantification Overview

After log2 value transforming and normalization, the datasets of quantified peptides with PTMs showed standard normal distribution (Figure 2a), validating the reliability of the data processing method. Principal component analysis (PCA) of peptides with PTMs showed a clear segregation of the three conditions, while samples from same time points but different replicates were clustered together (Figure 2b). The larger variability (52.3%) of component 1 was between stimulated samples (12h and 24h) and unstimulated samples (0h), indicating the fundamental effect of IFN-γ/TNF-α co-stimulation on the regulation of PTMs in INS-1E cells. The difference between 12h and 24h stimulation was separated in component 2 with smaller variability (19.2%). Very stringent criteria was applied for definition of significant regulation for proteins and peptides with PTMs, which should be observed in at least two replicates, with a p-value □ 0.05 in limma and rank tests, and showed no less than 1.5 fold change in one condition compared to the other conditions. The results revealed that IFN-γ/TNF-α co-stimulation caused significant regulation of 100 proteins at the expression level (Supplemental table S6), and much more regulations were observed at the PTM level among 3025 peptides with PTMs from 1490 proteins (Supplemental table S7). Multiple scatter plot analysis of regulated peptides with PTMs also revealed a high reproducibility between triplicates (Figure 2c), as the Pearson correlation values for same condition between triplicates were relatively high, varying between 0.81 and 0.93.

**Figure 2:**
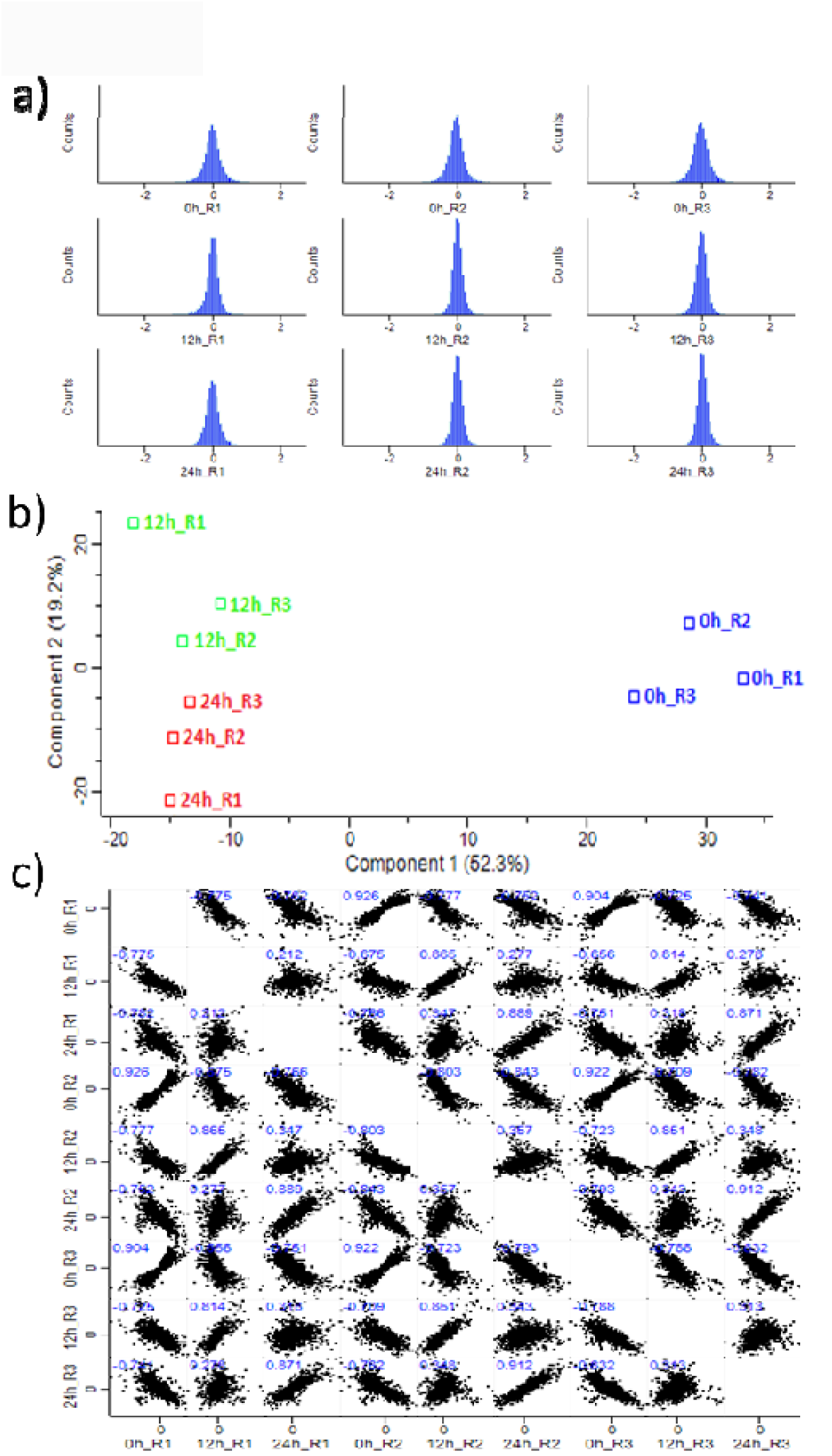
The statistical comparison of PTM data from INS-1E cells with cytokines stimulation at 0h, 12h and 24h in triplicates. (a) The normal distribution of PTM datasets. (b) The principle component analysis indicated the effects of cytokines stimulation on PTM changes at three time points. (c) The Pearson correlation analysis of PTM data in triplicate.

### Analysis of regulated proteins at the total protein level

For the 100 proteins with regulation at total protein level, compared to control (0h), after cytokines stimulation, 85 proteins showed increased expression, and only 15 proteins had decreased expression. Most of the upregulated proteins identified in this study were already confirmed to show up-regulated gene expression upon these cytokines stimulation using human cDNA microarray or oligonucleotide arrays [31, 32], and as so our results confirm their upregulation at the protein level. String network analysis (Figure 3a) revealed that the largest network was formed by the IFN-γ induced proteins, which all showed increased expression, and confirmed in the IFNs regulated genes database (http://www.interferome.org). The second largest network was clustered by the up-regulated immune proteins related to antigen processing and presentation, such as the major histocompatibility complex (MHC) antigens. The other three sub-networks included the up-regulated TNF-α induced proteins related to the NF-κB signaling pathway (CD40, NF-κB2 and ICAM1), proteasome proteins (PSME1, PSMB9 and PSMB10) and redox related proteins (SOD1, NOS2). The effect of cytokines on the upregulated expression of MHC antigens, CD40, NF-κB2 and ICAM1 were confirmed in a previous study [33]. NOS2 was observed with increased expression, and it’s well known that cytokines stimulation leads to increased NOS2 expression and NO production in β-cell [34]. SOD1, which is related to anti-oxidative stress in cells, was the only protein in the network with decreased expression after stimulation. Genetic ablation of SOD1 causes glucose intolerance, reduced in vivo beta cell insulin secretion and decreased beta cell volume [35], but overexpression of SOD1 can confer protection against toxicity of the cytokine mixture [36], Our results indicate that the cytokines stimulation may cause the oxidative stress through increased expression of NOS2 and decreased the amount of the antioxidant defense molecule SOD1. Furthermore, pathway analysis (IPA) revealed protein targets directly mediated by IFN-γ and TNF-α (Figure 3b); The network contained 36 proteins or protein complexes, among them, 35 and 25 targets were found to directly interact with IFN-γ and TNF-α respectively. Of these, 24 protein targets had interaction with both cytokines. This analysis indicated a highly shared interaction network map between IFN-γ and TNF-α.

**Figure 3:**
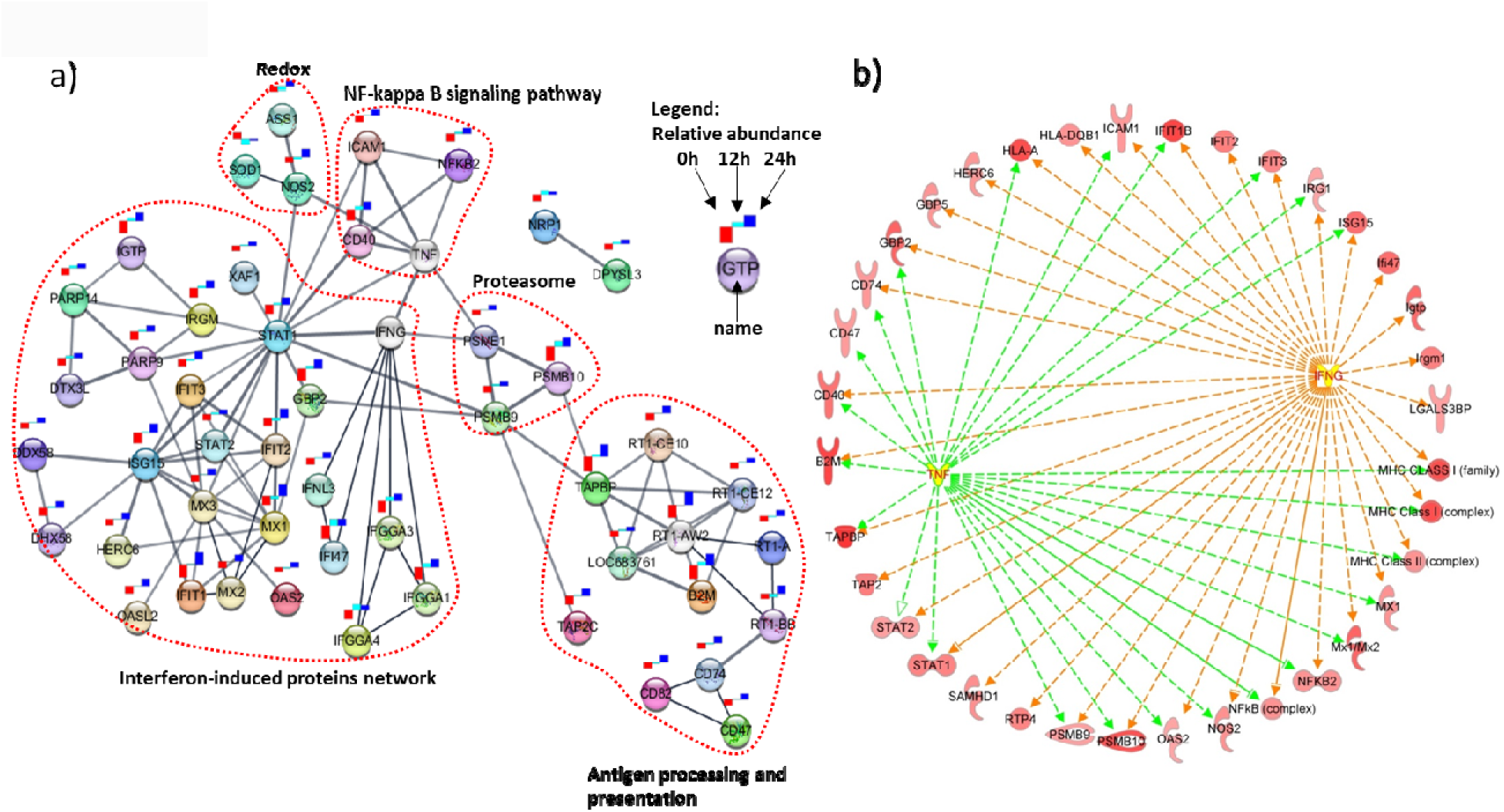
The protein-protein interaction network of proteins with regulation at total protein level. (a) The STRING protein-protein interaction network indicated the interested biological subnetwork with red dash line cycle, and the relative protein abundance at 0h, 12h and 24h was also indicated as annotated in legend. (b) IPA analysis revealed the IFN-γ and TNF-α directly interacting proteins, which were all up-regulated at the protein level after cytokines stimulation.

### Analysis of proteins with regulated PTMs

For the regulated peptides with PTMs, beside the previous mentioned statistical criteria, peptides with other possible induced modifications during sample preparation, such as unspecific deamidation on N and Q, oxidation on methionine were removed. The remaining peptides were manually inspected to confirm the reliability of the PTM sites. For the N-linked glycosites with NXS/T/C motif, the associated proteins must be secretory proteins or membrane associated proteins, on which N-linked glycosylation can happen. Based on these criteria, 3025 peptides with PTMs on 1490 proteins were found to be significantly regulated after stimulation (Supplemental table S7), as summarized in Supplemental Figure 2. A total of 1175 rmCys sites, 1514 phosphosites and 115 SAN-glycosylation sites were identified to be regulated in 809, 776 and 98 proteins respectively. String network analysis using Markov CLustering Algorithm (MCL) revealed more than 100 MCL protein sub-clusters, which should contain 3 or more proteins in a cluster (Supplemental Figure 3 and Supplemental table S8). The analysis revealed that these three PTMs demonstrated specific spatial and biological processes related distribution patterns. Generally, protein phosphorylation and reversible Cys modifications were the two dominant PTMs, widely distributed and regulated in most of the listed proteins, but they also showed differential distributions for specific protein clusters. N-linked glycosylation was rarely observed in a few protein clusters. The most intense protein clusters for each PTM were presented in Figure 4a, b, c. The rmCys modification was highly represented in networks related to protein translation in ribosome and proteolysis in proteasome, RNA binding and ribosome biogenesis, glycolysis and glucogenesis, etc. (Figure 4a). Phosphorylation was the main regulated PTM in networks related to mitotic cell cycle, RNA splicing in spliceosome, small GTPase signaling, transmembrane signaling, etc. (Figure 4b). Regulated SAN-glycosylation was relatively over-represented in networks related to MHC RT1 class proteins, antigen processing and presentation proteins, immune response related complements (Figure 4c). The most intense protein cluster with 45 proteins was related to protein translation in ribosome and proteolysis in proteasome, and most proteins in this cluster were found with only regulated rmCys. IPA analysis of regulated proteins revealed a more complicated network at the PTM level, including 79 proteins that directly interact with INFγ and TNFα (Figure 5), the proteins in the outer ring of the network were interacting only with TNF-α (connected with pink line) or IFN-γ (connected with orange line), whereas the 43 proteins in the inner rings were interacting with both cytokines, and some of the interesting key proteins were indicated in the most inner ring of the network, such as STATs, NOS2, SODs, NF-κB complex, ICAM1 and MXs. Whereas many of the proteins in these networks showed regulated phosphorylation, more proteins contained regulated Cys modification. Regulated N-linked glycosylation was identified only on 10 proteins in the network, which were all membrane associated proteins.

**Figure 4:**
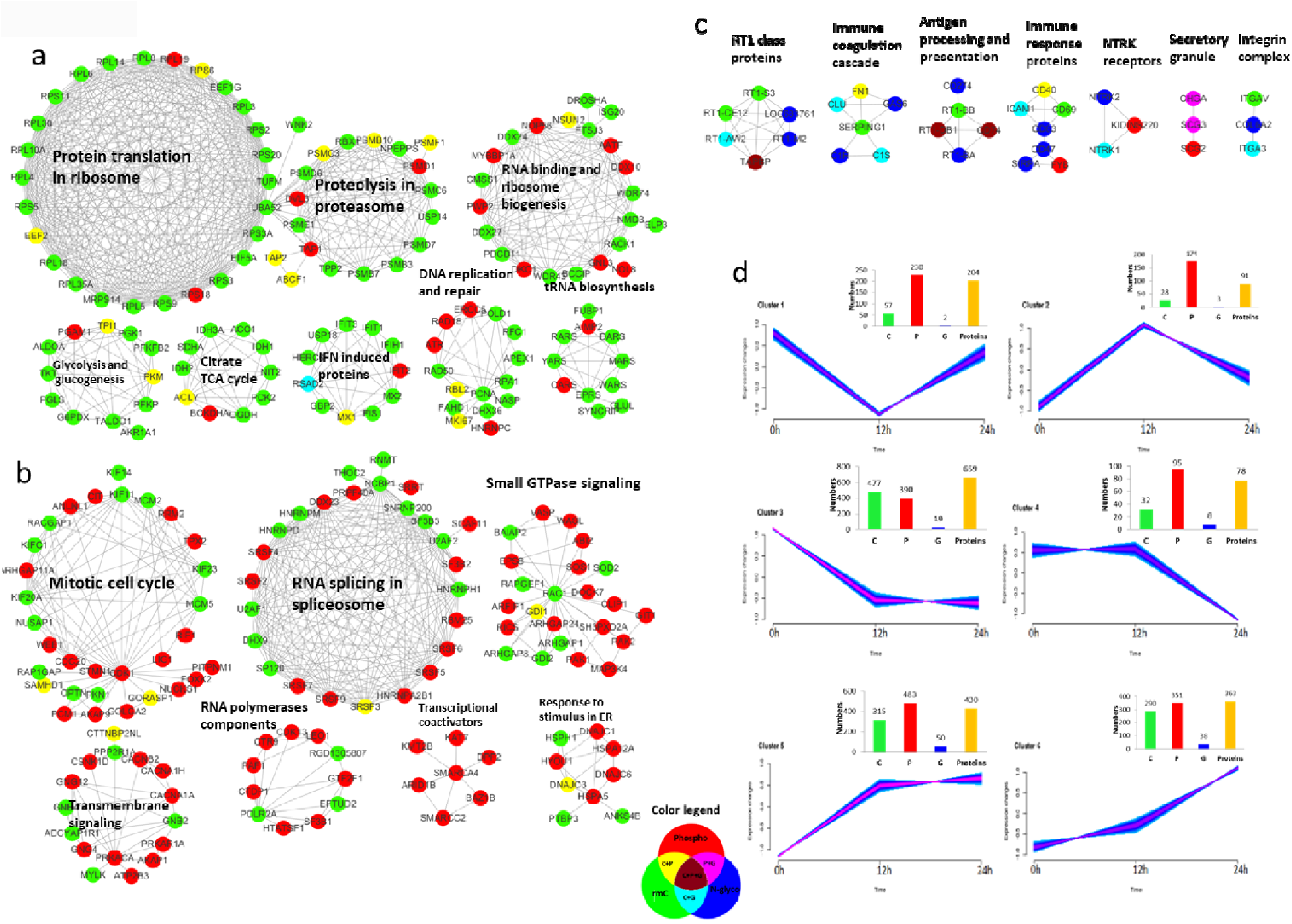
The network and cluster analysis of proteins with regulated PTMs revealed that the three PTMs demonstrated specific temporal patterns related to diverse biological processes in the time course of cytokine stimulation. The top seven most intense protein clusters for each PTM from STRING network analysis were presented in (a) for rmCys, (b) for phosphorylation and (c) for SAN-glycosylation. The color indicated the type of regulated PTMs in the protein as presented in the legend. (d) The fuzzy c-means clustering analysis revealed different temporal change patterns of three PTMs during cytokines stimulation from 0 h to 24 h, the numbers of related PTM sites and proteins were also presented for each cluster.

**Figure 5:**
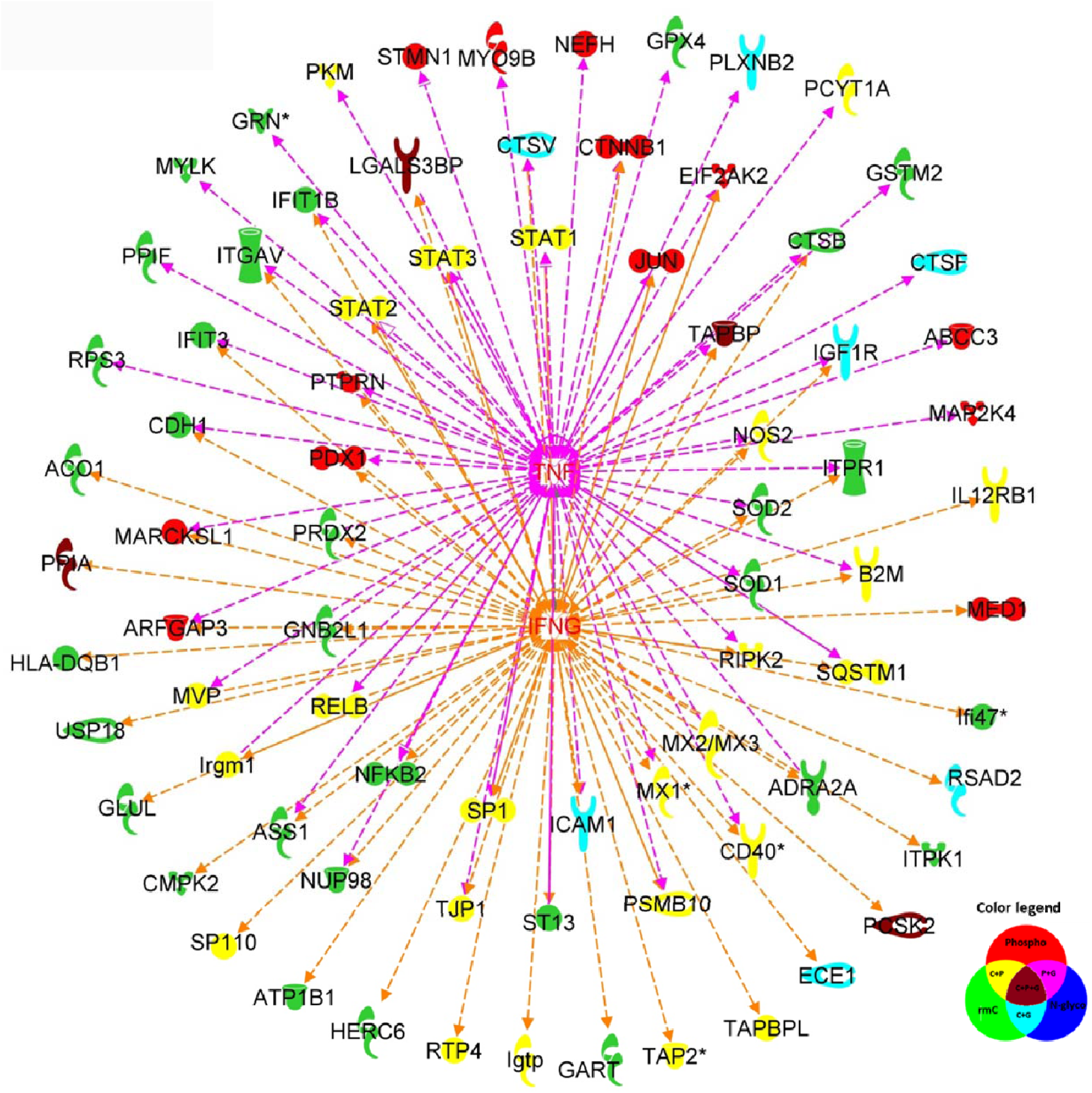
IPA analysis revealed the IFN-γ and TNF-α directly interacting proteins with regulated PTM sites. The color indicated the type of regulated PTMs in the protein.

### Cluster analysis revealed different spatial and temporal patterns of the three PTMs

The fuzzy c-means clustering method was used to analyze the time-course change patterns of regulated peptides with PTMs, and revealed 6 clusters (Figure 4d and Supplemental table S7). The numbers of regulated sites for the different PTMs and proteins in each cluster were also indicated on the right-up corner of each cluster. The PTM sites in cluster 1 and cluster 2 showed a quick response to cytokines stimulation, the effects were maximized at 12h (decreased in cluster 1, increased in cluster 2), and then relatively attenuated to almost normal level at 24h. Phosphorylation was the main regulated PTM in cluster 1 and 2. The PTM sites in cluster 3 and cluster 4 generally showed a down-regulated pattern after cytokines stimulation, and the majority of them were in cluster 3 with decrease to a low level and stable from 12h to 24h. More rmCys sites were identified than other PTM sites in cluster 3, indicating decreased level of reversible Cys modification. Cluster 5 and cluster 6 represented PTMs that were up-regulated after stimulation. More regulated phosphosites were observed than regulated rmCys sites. Interestingly, much more up-regulated N-glycosylation sites were identified in cluster 5 and 6 than in other clusters, and highly represented in immune related networks, reflecting the important role of N-glycosylation in cytokines mediated immune response. The analysis indicated the three regulated PTMs also demonstrated different temporal patterns in the time course of cytokine stimulation, indicating different regulatory roles of these PTMs.

String network analysis of proteins in each cluster further revealed the associated biological processes for the regulated PTMs. Proteins with regulated PTMs in cluster 1 and 2 (Supplemental Figure 4) were related to translation, RNA splicing in spliceosome, metabolism processes, and some networks centered to specific proteins such as CDK1 and RAC1. Phosphorylation was the dominant regulated PTM for these proteins. Many more proteins were present in cluster 3, these proteins were mainly regulated by rmCys, and they were related to series of biological processes (Supplemental Figure 5), such as DNA replication and repair, RNA transcription, splicing, and transport, protein translation and processing, citrate/TCA cycle, carbohydrate and purine metabolism etc. No complicated network was observed in cluster 4. Proteins in cluster 5 and 6 (Supplemental Figure 6) were related to networks involved in DNA, RNA, protein and metabolism processes. Other very interesting networks were also enriched in these two clusters, including proteasome proteins, cytokines induced proteins and related interactions, cytokines regulated signaling pathways, such as JAK-STAT signaling, NF-κB signaling, and a large number of immune proteins related to antigen processing and presentation. Essentially, up-regulated N-glycosylation was found to be highly represented in these immune related networks, reflecting the important role of N-glycosylation in cytokines mediated immune response. Regulated phosphorylation and rmCys were both widely distributed in cluster 5 and 6. Interestingly, phosphorylation and rmCys were both found to be regulated in proteins related to RNA splicing in the spliceosome. However, these two PTMs showed different regulation patterns, proteins with regulated phosphorylation were observed in cluster 1, 2, 5 and 6, but proteins with rmCys were predominantly found in cluster 3, indicating different regulatory roles for phosphorylation and rmCys. In summary, the combined cluster and network analysis indicated that the rmCys, phosphorylation and N-glycosylation were generally regulated by pro-inflammatory cytokines and widely distributed in INS-1E cells. However, they presented specific spatial and temporal patterns related to diverse biological processes in the time course of cytokine stimulation.

**Figure 6:**
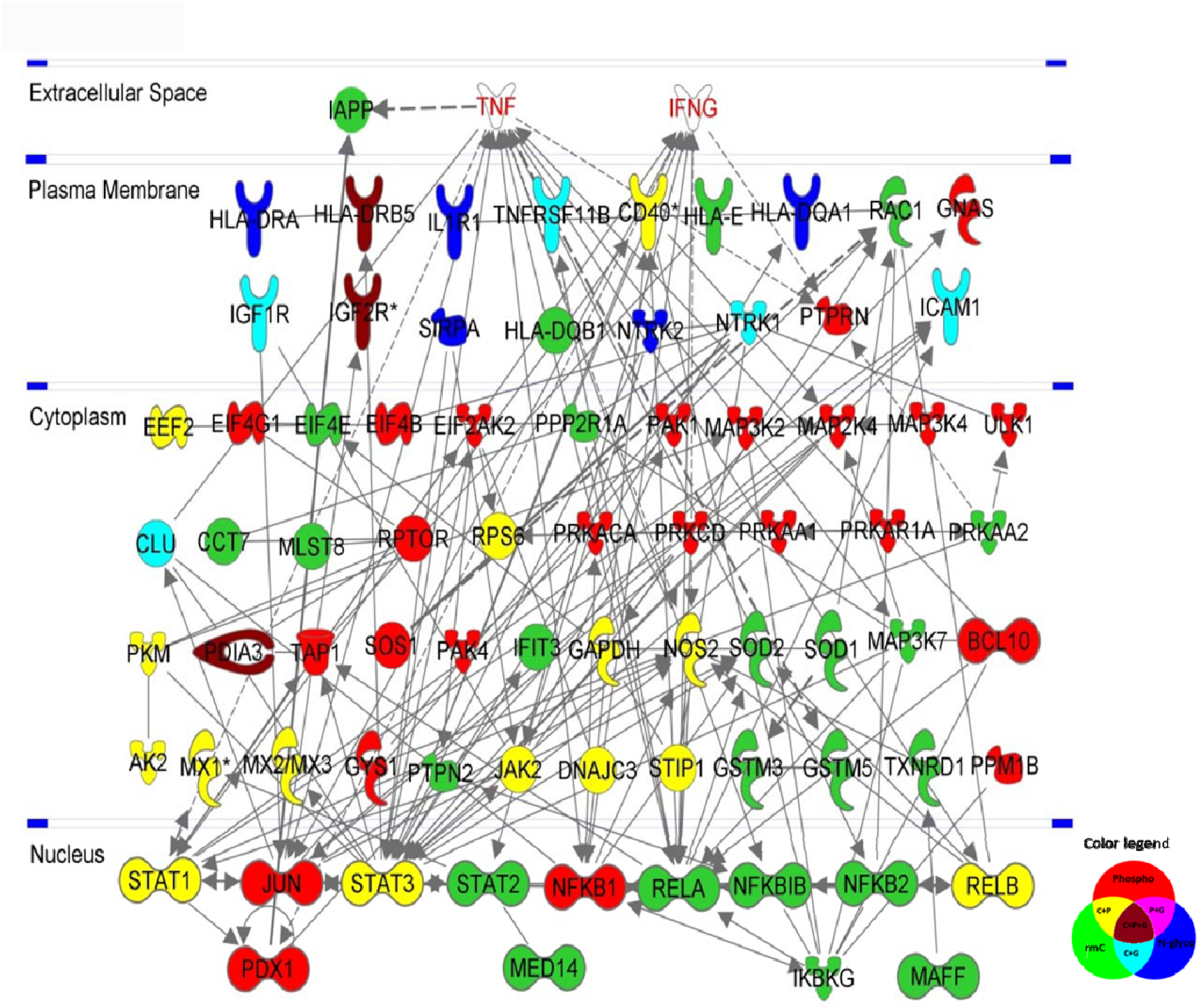
The protein interaction network of proteins with regulated PTMs. It was built based on interested signaling pathways using IPA analysis. The color indicated the type of regulated PTMs in the protein as presented in legend.

### Signaling pathway and interaction analysis

IPA signaling pathway analysis was further employed for signaling pathways and interaction analysis of the proteins with regulated PTMs, it revealed 16 interesting over-represented signaling pathways (Supplemental table S9). These pathways were cytokines and cytokines mediated pathways, and pathways related to diabetes, redox regulation, immunological and metabolic activities. The proteins with regulated PTMs in these selected pathways were used to build the interaction network. It was built using data source from specific tissue and cell lines (the pancreas, beta cell, and pancreatic cancer cell lines) at high confidence level in order to increase the specificity and decrease the complexity. An interesting interaction network (Figure 6) was revealed and presented based on the interaction and cellular location information of these proteins. The proteins were differentially colored to indicate their regulated PTMs. The network demonstrated that TNF-α and IFN-γ co-stimulation led to the regulation of the three PTMs in proteins from membrane to nucleus, rmCys and phosphorylation were regulated in proteins located in whole cell, whereas N-linked glycosylation were regulated mainly in membrane proteins, which was in agreement with the property of N-linked glycosylation. The pro-inflammatory cytokines TNF-α and IFN-γ initiate the activation of NF-κB, STATs and mitogen-activated protein kinases (MAPK) as well as their related β-cell gene networks, leading to the activation of NOS2 and increases in nitric oxide (NO), which ultimately induce β-cell apoptosis [21, 34]. Our data confirmed that cytokines co-stimulation induced the upregulation of STATs, NF-κB and NOS2 at total protein level(Figure 3a). The IPA analysis further revealed the different regulation of PTMs in these key proteins and related proteins in the network (Figure 6). As indicated, the STAT signaling molecules: JUN and STATs, the NF-κB complex proteins: NF-κB1, NF-κBIB, NF-κB2, REL-associated proteins RELA, RELB and inhibitor of NF-κB kinase subunit gamma (IKBKG) and pancreatic and duodenal homeobox 1 (PDX1) essential for beta cell function were located in the nucleus. STATs and RELB were regulated by both phosphorylation and rmCys, but RELA, NF-κBIB, NF-κB2 and IKBKG were identified with only regulated rmCys, JUN, PDX1 and NF-κB1 were identified with only regulated phosphorylation. In the cytoplasm, NOS2 was identified with both regulated phosphorylation and rmCys, JAK1 and MAPKs were identified with only regulated phosphorylation. On the membrane, regulated N-linked glycosylation and other PTMs were mainly identified on protein receptors, ICAM1 and immune proteins related to antigen processing and presentation.

### Characterization of regulated signaling pathways related to NOS2

The activation of transcriptional factors JUN, STAT1 and NF-κB by cytokines can induce the expression of NOS2, lead to the increased production of NO, and mediate the apoptosis of β-cells [21], as indicated in the NOS2 signaling pathway (Figure 7). In our study, the increased expression and regulated phosphorylation and rmCys were observed on STATs upon cytokine stimulation. Phosphorylation of STAT1 induced by IFN-γ was reported to regulate the STAT1 transcriptional activity [37]. We observed that the phosphorylation of S523 on JAK2 was upregulated after stimulation, and it can serve as a negative regulator to dampen activation of JAK2 in response to stimulation [38]. Phosphorylation of S63 and S73 on transcription factor AP1 (JUN) was also identified to be upregulated to high level at 12h and then decreased at 24h of stimulation. The phosphorylation of S63 and S73 can induce the activation of JUN and regulate cell cycle regression and apoptosis [39]. The NF-κB complex has five members, NF-κB1, NF-κB2, RELA, RELB and c-REL. Activation of the NF-κB is mediated by the IκB kinase (IKK), and the inhibitor of nuclear factor kappa-B kinase subunit gamma (IKBKG or IKK-γ) is a “master” regulatory protein subunit of IKK [40]. In our study, the NF-κB2 was highly increased at the protein level after stimulation, meanwhile, increased phosphorylation was observed on S493 in NF-κB1 and S549 in RELB. Multiple regulated rmCys sites were also observed on different protein subunits of the NF-κB complex (Figure 7). The C340 in rat IKBKG, which is identical to C347 of human IKBKG, can form intermolecular disulfide bond in the IKBKG dimer with C54, which is mediated by ROS and facilitate the activation of NF-κB DNA binding in response to treatment with TNF-α [40]. The observed upregulation of this site in our study may indicate the formation of more dimers to enhance the NF-κB DNA binding. The increased expression and up-regulated rmCys and phosphorylation of NOS2 were also identified. C107 and C112 in rat NOS2 are identical to C104 and C109 of human NOS2, which can contribute to form the zinc tetrathiolate cluster to maintain the stability and activity of the homodimeric NOS2, as monomeric NOS2 is inactive [41]. The increased rmCys of C107 and C112 in NOS2 may increase NOS2 activity and NO production through this mechanism.

**Figure 7:**
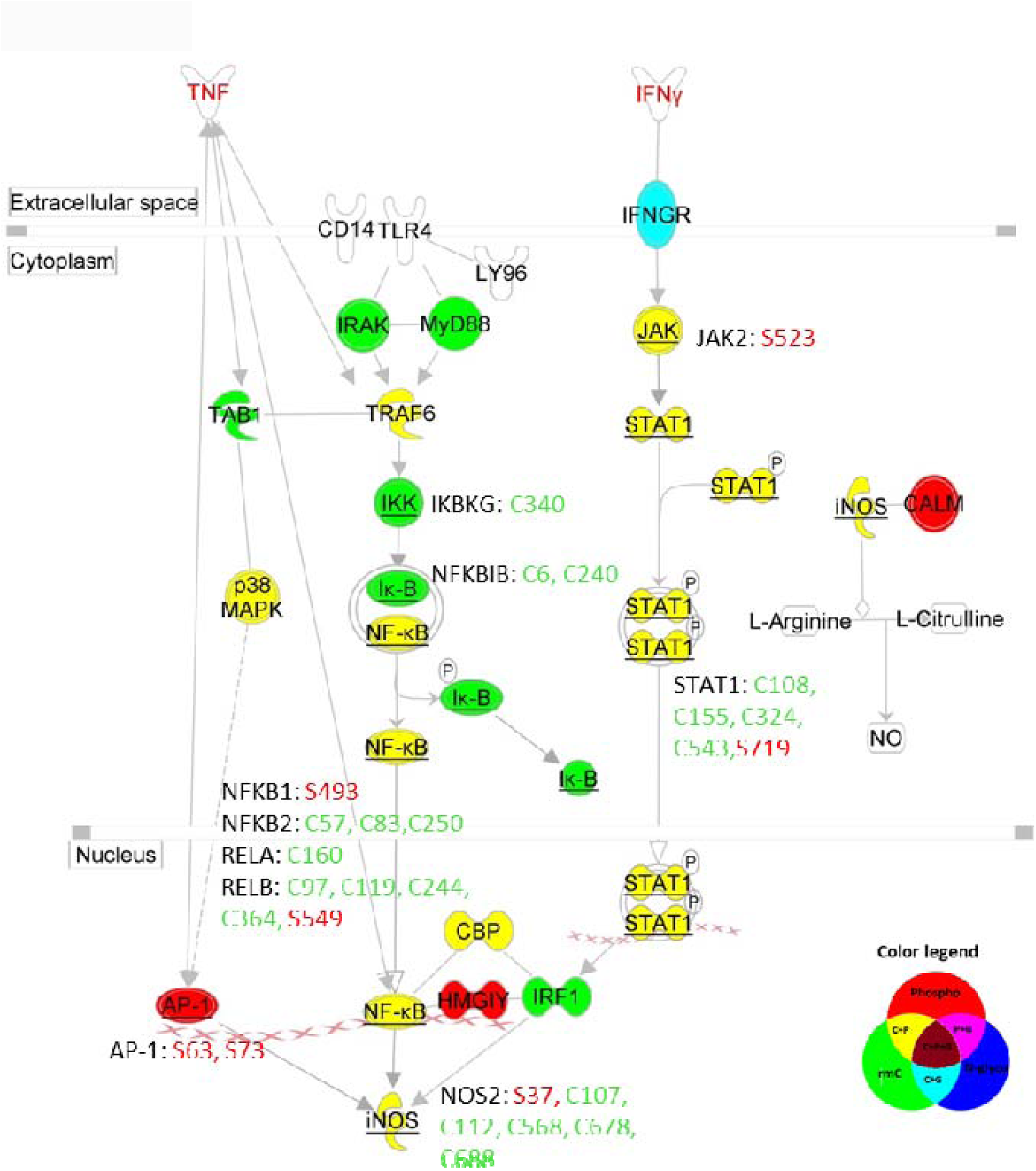
The NOS2 signaling pathway. The proteins identified with PTMs were labelled with different colors in the pathway, and for the proteins with the regulated PTM sites, the sites were also indicated. The color indicated the type of regulated PTMs in the protein as presented in legend.

Beside the molecules related to NOS2 signaling, other essential proteins related to redox and beta cell function were also identified with regulated PTMs, such as SOD1 and SOD2. They are antioxidant enzymes that catalyze the dismutation (or partitioning) of the superoxide (O_2_^−^) radical into either ordinary molecular oxygen (O_2_) or hydrogen peroxide (H_2_O_2_). SOD1 is located in the cytoplasm, and SOD2 is located in the mitochondria. We observed that the expression and the rmCys site C7 of SOD1 were significantly decreased after 12h cytokines stimulation, and increased a bit after 24h cytokines stimulation. This Cys site in human was identified to be a palmitoylation site, and mutation of this site prevented palmitoylation, leading to reduction of SOD1 activity in vivo and in vitro, and inhibition of its nuclear localization [42]. The decreased expression and Cys modification may indicate a reduced anti-oxidant activity in the cell upon cytokine stimulation. C220 in SOD2 was identified to be significantly upregulated, in contrast to SOD2 protein expression after stimulation. Heterozygous SOD2 deletion in mice, a model that mimics SOD2 changes observed in diabetic humans, can impair glucose-stimulated insulin secretion, in high fat-fed mice without affecting insulin action [43]. The increased Cys modification might be a protective mechanism in mitochondria against oxidative stress caused by cytokines stimulation.

### N-linked glycoproteins related to receptors and immune response

Compare to Cys modification and phosphorylation, N-linked glycosylation was much rarely observed, and its regulation was found mostly on proteins associated with the membrane, and these proteins mostly were receptors and proteins related to immune response. For receptors, upregulated N-linked glycosylation and Cys modification were identified on N901, C215, C219, C332, C456, C489, C664, C807 and C816 in Insulin-like growth factor 1 receptor (IGF1R), while IGF2R was identified to be downregulated at total protein level, and multiple PTM sites were also found to be downregulated after stimulation, including N737, N2125, C64, C366, C375, C883, C1259, C1450, C1966, C2028, C2114, S2397, S2467, S2472. According to the Uniprot database, these regulated PTM sites were identified in human IGF1R and IGF2R by large scale MS analysis as well, and most of the Cys sites were responsible for the formation of disulfide bonds and related to the dimerization and activation of receptors [44]. Adequate N-linked glycosylation is required for the translocation of IGF receptors to the cell surface, and inhibition of N-linked glycosylation resulted in down-regulation of IGF-1R at the cell surface, and correlated with a drastic decrease in IGF-1R auto-phosphorylation and inactivation of the receptor [45]. However, this study doesn’t report on the altered PTM sites, which we observed here. TNFRSF11B and CD40 are TNF receptors, the PTM sites C124, C185, N178 and N289 in TNFRSF11B, S49, C51, C83, C186 and C259 in CD40 were identified to be upregulated after stimulation, human studies reported that most of these Cys sites can form disulfide bonds essential for the dimerization of CD40 [46] and trimerization of TNFRSF11B [47]. The two glycosylation sites of TNFRSF11B were also reported in Uniprot database. In addition, N67 and C659 of high affinity nerve growth factor receptor 1 (NTRK1) were found to be downregulated and N205 of NTRK2 were upregulated, and these glycosylation sites were also confirmed by other study in Uniprot database [48]. Multiple proteins related to immune response were identified with regulated sialylated N-linked glycosites, including ICAM1 and several major histocompatibility complex (MHC) proteins encoded by the human leukocyte antigen (HLA) gene complex. ICAM1 is a transmembrane protein of the immunoglobulin superfamily, and the ligand for the leukocyte adhesion protein integrin αLβ2. Its expression can be significantly increased in the presence of cytokines and ROS [49]. ICAM1 is characterized as heavily glycosylated, and N-glycosylation deficiency would reduce ICAM1 induction and impair inflammatory response [50, 51]. The increased expression of ICAM1 was also observed in our study, meanwhile, up regulation of PTMs was determined on 4 N-linked glycosylation sites (N154, N202, N309, N464) and 3 rmCys sites (C135, C290, C382) of ICAM1 after normalization to the protein level. These regulated PTM sites were already confirmed in structural studies and annotated in the Uniprot database. The regulated Cys sites can form disulfide bonds with other Cys sites in the structure of ICAM1 and they are essential for the affinity with integrin αLβ2 and dimerization [52, 53]. The upregulation of PTM on these sites should be triggered by the immune response induced by cytokines. All MHC family members carry N-linked glycosylation at sites that are highly conserved across evolution, N-linked glycosylation is essential for protein structure and quality control events in the ER and additional functional roles such as mediating receptor–ligand interactions and/or molecular geometric spacing for immune responses [54]. In our study, multiple MHC proteins were upregulated at the total protein level, as indicated in Figure 3b, meanwhile, N105 in HLA-DQA1, N96 in HLA-DRA and N46 in HLA-DRB5 were identified with upregulated N-linked glycosylation after normalization (N105 of HLA-DQA1 is a conserved site identical to N96 of HLA-DRA). These data strongly indicate that sialylation is heavily involved in these immune responses to cytokine stimulation.

Additionally, multiple modified Cys sites and phosphosites on MHC proteins were also found to be upregulated after normalization. Mx proteins are interferon-induced members of the dynamin superfamily of large GTPases with essential function in early antiviral host defense [55]. In this study, MX1, MX2 and MX3 were all found to be upregulated at the total protein level upon cytokines stimulation, in agreement with another study [56]. At the PTM level, the modified PTM sites C33, C313, C327, C578 and S557 in MX1, C40 in MX2, C40 and S190 in MX3 were mostly found to be up-regulated even after normalization to the protein abundance. C40 is conserved in MX2 and MX3. However, the function of these PTM sites were not annotated, it would be a very challenging task to characterize the function of regulated PTM sites identified in this study.

## Conclusion

Combined analysis of multiple PTMs in an easy and efficient manner is still a very challenging task, but highly desired in MS based PTMomics studies. In this work, we developed the ‘one stone for three birds’ comprehensive TiCPG strategy for multiple PTMs. The TiCPG strategy enables the simultaneous characterization of the proteome and three essential PTMomes with high efficiency and specificity using low amount of starting material, and it also greatly minimizes the sample loss problem observed in other strategies targeting multiple PTMs. The complete procedure is relatively straight-forward and easy to perform in most laboratories, with one step enrichment for peptides with three PTMs and subsequent separation and pre-fractionation steps. The identification coverage for each PTM using the TiCPG strategy is comparable to that of individual PTM analysis. In principle, the TiCPG strategy can also be extended to include other PTMs that can be enriched by TiO_2_ chromatography, and it can also be integrated with other enrichment strategy to include additional PTMs, such as antibody-based enrichment of ubiquitinated and lysine acetylated peptides. The application of the TiCPG strategy to characterize the proteome and the targeted three PTMs in INS-1E cells subject to IFN-γ/TNF-α co-stimulation revealed differential spatial and temporal patterns related to protein function and cellular localization in the time course of cytokine stimulation. We believe that the TiCPG strategy will be a powerful tool for characterization of multiple PTMs, PTM cross-talk and related interactions in biological or clinical applications.

## Materials and Methods

### Materials

All chemicals were purchased from Sigma-Aldrich (St. Louis, MO), unless otherwise stated. Modified trypsin was from Promega (Madison, WI). Poros R2 and Poros Oligo R3 reversed-phase material were from Applied Biosystems (Forster city, CA). GELoader tips were from Eppendorf (Hamburg, Germany). The 3M Empore™ C8 disk was from 3M Bioanalytical Technologies (St. Paul, MN). Titanium dioxide beads were a gentle gift from GL Sciences Inc. (Tokyo, Japan). N-Succinimidyl Iodoacetate (SIA) was purchased from Thermo Scientific (Rockford, IL). PNGase F was obtained from New England Biolabs (Ipswich,MA). Glyko® Sialidase C™ was from Prozyme (Hayward, CA). All solutions were made with ultrapure Milli-Q water (Millipore, Bedford, MA).

### Cell culture

Rat insulinoma INS-1E cells were cultured in RPMI 1640 GlutaMAX medium (Invitrogen # 61870) supplemented with 5% FBS (Biochrom, # S 0115), 1% Penicilin/streptomycin (Invitrogen # 15070-063), 1mM sodium pyruvate (sigma Aldrich #S8636) and 50 µM 2-mercaptoethanol in 10 cm falcon DB plates (#353003). Cells were split with trypsin and maintained at 37°C in a humidified atmosphere containing 5% CO2.

At 90% confluency the cells were aspirated in serum free RPMI 1640 GlutaMAX medium with 1% Penicilin/streptomycin (Invitrogen # 15070-063), 1mM sodium pyruvate (sigma Aldrich #S8636) and 50 µM 2-mercaptoethanol) for 10 hours. The cells were subject to cytokines stimulation in serum free medium containing 10 ng/ml rat IFN-γ (Peprotech, #400-20) and 10 ng/ml rat TNF-α (Peprotech, 400-14) for 12 and 24 hours. The control cells were aspirated in new serum free medium supplemented with penicilin/streptomycin, sodium pyruvate and 2-mercaptoethanol for 5 minutes before harvesting. After stimulation the cells were washed in cold 1x PBS, and harvested by scraping, centrifuged at 300 g for 3 minutes, PBS removed and snap frozen in liquid nitrogen. The cell pellets were stored at −80°C until further used.

### Preparation of protein lysate from cytokines stimulated INS-1E cells

Triplicates of the control INS-1E cells, and the cells subject to 12h, 24h IFN-γ/TNF-α stimulation from 15cm dishes respectively were re-suspended in 500 μL lysis buffer containing 6 M urea, 2 M thiourea, 2% SDS, 40 mM N-Ethylmaleimide (NEM), complete protease inhibitor (Roche, Hvidovre, Denmark, one tablet per 50 mL) and phosphatase inhibitor PhosStop (Roche, Hvidovre, Denmark, two tablets per 50 mL), and lysed on ice using a probe tip sonicator. After using 10KDa spin filters to remove SDS and excess of NEM, The protein solution in the filter was diluted in 500 μl 50 mM TEAB buffer, and the protein concentration was determined by Qubit assay (Invitrogen, Waltham, MA). Then the proteins were reduced with 10 mM tris (2-carboxyethyl) phosphine (TCEP) for 1h at room temperature. No alkylation step was performed here. The reduced proteins were subsequently digested with 0.04 AU Lys-C (Wako, Japan) for 3h at room temperature, and trypsin (2%, w/w) was added for further digestion at 37°C overnight. The samples were acidified to a final concentration of 2% formic acid and 0.1% trifluoroacetic acid (TFA) and centrifuged at 20000 × g for 30 mins to precipitate the lipids. The acidified peptides were desalted and purified with a Hydrophilic-Lipophilic-Balance solid phase extraction (HLB-SPE) (Waters, Bedford, MA) cartridge according to the manufacturer’s instructions, and the eluted peptides from HLB column were lyophilized and subsequently resuspended in 200 μL 50 mM TEAB buffer before isobaric tagging for relative and absolute quantification (iTRAQ) labeling, and the concentration was measured using Qubit assay again. A total of 100 μg peptides from each time course was labeled with iTRAQ™ (Applied Biosystems, Foster City, CA) as described by the manufacturer (0 hour/control, iTRAQ-114; 12 hours, iTRAQ-115; 24 hours, iTRAQ-116). After labeling, the samples were mixed 1:1:1 and dried by vacuum centrifugation. All experiment was performed in biological triplicate.

### Alkylation with synthesized CysPAT at peptide level

The dried iTRAQ labelled peptides mixture was dissolved in 200 μl 0.1% TFA buffer, desalted and purified with in-house prepared Oligo R3 Reversed phase column, eluted from the R3 column with 70% acetonitrile (ACN), 0.1% TFA and dried by vacuum centrifugation. The CysPAT tag was synthesized as previously described [25, 26]. The purified iTRAQ labelled peptides were resuspended and mixed with 10mM of the synthesized CysPAT and 5 mM TCEP in 50 mM TEAB, pH 7.8 in a volume of 200 µl. The mixed solution was incubated in the dark for 1h with gentle agitation. Afterwards, the solution was subjected to Oligo R3 reversed phase column purification to purify the peptides and remove extra chemicals. The eluted peptides were lyophilized.

### TiO_2_ enrichment

TiO_2_ enrichment was performed to simultaneously enrich the phosphopeptides, CysPAT labelled reversibly modified Cys (rmCys) peptides and sialylated N-linked (SAN-) glycopeptides as previously described for phosphorylated peptides [23, 25, 28]. A total of 1.8 mg TiO_2_ per 100 μg peptides was added to ensure efficient enrichment of peptides with these three PTMs.

The labelled peptides was dissolved in 100 μl 0.1% TFA, and then diluted 10 times with loading buffer of 80% ACN, 5% TFA and 1 M glycolic acid. A total of 5.4 mg TiO_2_ (1.8 mg TiO_2_ per 100 μg peptides) was added to the solution, shaken for 10 min at 600 rpm and centrifuged. The supernatant was collected carefully and incubated with half the amount of TiO_2_. The TiO_2_ beads were firstly washed with 100 μL of loading buffer by mixing for 15 s, transferred to a new tube and centrifuged to pellet the beads, then washed with 100 μL of a solution containing 80% ACN and 1% TFA, and followed by washing with 100 μL of a solution containing 20% ACN and 0.1% TFA. The peptides were eluted with 100 μL elution buffer (40 μL 28% ammonia solution in 960 μL water, pH 11.3) by shaken for 10 min and centrifuged for 1 min. The eluent was collected and passed through a C8 stage tip to remove TiO_2_ beads and the peptides attached to the C8 tip were subsequently eluted with 10 μL 30% ACN. The eluted peptides were dried by vacuum centrifugation to obtain the enriched multiple PTMs peptide fraction containing rmCys peptides, phosphopeptide and SAN-glycopeptides. The unbound peptides and subsequent washes were combined and dried by vacuum centrifugation to obtain the non-modified peptide fraction. The non-modified peptide fraction was resuspended in 0.1% TFA, purified by an Oligo R3 micro-column and dried by vacuum centrifugation.

### Deglycosylation and second TiO_2_ enrichment

The enriched multiple PTMs peptide fraction was resuspended in 100 μl of 50 mM TEAB, pH 8.0 and deglycosylated with 500 U of PNGase F (New England Biolabs, Ipswich, MA) and 0.1 U Sialidase A (Prozyme, Hayward, CA) for 12 h at 37 °C [57]. After incubation, the peptide mixture was acidified by 1% TFA and dried by vacuum centrifugation. Then the peptides were dissolved in 100 μl buffer of 0.1% TFA and 70% CAN, and subject to second round TiO_2_ enrichment. The eluted peptides from the second TiO_2_ enrichment were acidified and dried by vacuum centrifugation to produce the co-enriched rmCys peptides and phosphopeptides fraction. The unbound peptides and subsequent washes were pooled and dried by vacuum centrifugation to obtain the deglycosylated formerly SAN-glycopeptide fraction. Finally, both dried PTM peptide fractions were resuspended in 0.1% TFA, purified by Oligo R3 micro-columns and dried by vacuum centrifugation for further pre-fractionation.

### Hydrophilic Interaction Liquid Interaction Chromatography (HILIC)

After the above enrichment steps, we got three fractions of peptides, the non-modified peptide fraction, the co-enriched rmCys peptides with CysPAT labeling and phosphopeptides fraction and the deglycosylated formerly SAN-glycopeptide fraction. All these three peptide fractions were further fractionated using HILIC. The peptides were fractionated on an in-house packed TSKgel Amide-80 HILIC (Tosoh Bioscience, 5 μm) 320 μm × 170 mm μHPLC column by using the Agilent 1200 micro-HPLC instrument[57]. Briefly, the samples were suspended in solvent B (90% ACN, 0.1% TFA) by adding 10% TFA followed by water and finally the acetonitrile was slowly added to the aqueous solution in order to prevent peptide precipitation by the high acetonitrile concentration. Peptides (no more than 40 μg) were loaded onto a 320 ID peak HILIC column and eluted at 6 μL/min by decreasing the solvent B concentration (100–60% ACN, 0.1% TFA in water) in 42 min. Fractions were automatically collected in a 96 well plate at 1 min intervals after UV detection at 210 nm and dried by vacuum centrifugation, and stored at −20 °C until LC–MS/MS analysis.

### Mass spectrometric detection

After HILIC, all the pre-fractionated peptides were analyzed with a nanoEasy LC (Thermo Scientific, Odense, Denmark) combined with an Orbitrap Fusion Tribrid™ Mass Spectrometer (Thermo Fisher Scientific). HILIC fractions were resuspended in 5 µL of buffer A (0.1% FA), loaded onto a 2 cm 100 ID C18 pre-column with buffer A using a nanoEasy LC and separated on an in-house packed Reprosil-Pur C18-AQ (3 µm; Dr. Maisch GmbH, Germany) analytical column (20 cm x 75 µm ID). The gradients were set as: 0% – 34% buffer B (90% Acetonitrile, 0.1% FA) in 90 min, 34% – 100% buffer B in 5 min, and 100% buffer B in 8 min at a flow rate of 250 nL/min. Eluted peptides were analyzed on an Orbitrap Fusion mass spectrometer. Survey scans of peptide precursors from 400 to 1400 m/z were performed at 120000 resolution (at 200 m/z) with a 4 × 10e5 ion count target, the max injection time was 50 ms. Tandem MS was performed by isolation at 1,6 Th with the quadrupole, HCD fragmentation with normalized collision energy of 35, and MS2 analysis in the Orbitrap. The MS2 resolution was set to 30,000 and the max injection time was 100 ms. Only those precursors with charge state +2, +3 and +4 were sampled for MS2. The dynamic exclusion duration was set to 20 s with a 10 ppm tolerance around the selected precursor. The instrument was run in top speed mode with 3s cycles, meaning the instrument would continuously perform MS2 events until the list of non-excluded precursors diminishes to zero or 3 s, whichever is shorter. A fixed first mass of 100 was used. Raw data were submitted to pride (http://www.ebi.ac.uk/pride/archive/) under the project accession PXD007173 (username, password: 0Vci1jNE).

### Database searching, statistical and bioinformatics analysis

The LC-MS/MS data were processed with Proteome Discoverer (Version 1.4.1.14, Thermo Fisher Scientific) and subjected to database searching using both an in-house Mascot server (Version 2.2.04, Matrix Science Ltd., London, UK) and the embedded Sequest HT server with the following criteria: database, SwissProt Rattus norvegicus (Rat) and Mus musculus (mouse) protein database (updated to 15-04-2016); enzyme, trypsin; maximum missed cleavages, 2; variable modifications included oxidation (Met), acetyl (protein N-terminus and lysine(K)), SIA for Cys (SIA represented the CysPAT tag), NEM for Cys, deamidation for Asn and Gln, phosphorylation (Ser, Thr, and Tyr), iTRAQ were also included as a variable modification. The MS and MS/MS results were searched with a precursor mass tolerance at 10 ppm and a MS/MS mass tolerance at 0.05 Da. The results were filtered in Proteome Discoverer with the integrated Percolator algorithm [58] to ensure the false discovery rate (FDR) less than 0.01. Only peptides identified with high confidence (unambiguous or selected), first rank, Mascot ion score higher than 18 and passed the default score versus charge state for Sequest HT were accepted. For phosphorylation sites, the related PhosphoRS probability should be no less than 95% [59]. Only potential N-linked glycosylation sites with deamidation of asparagine (N) within the eukaryotic glycosylation sequon (NXS/T/C, X ≠ P) were considered. Peptides with different amino acid sequences or modifications were considered unique. The generated quantitative data was further filtered by removing the data with missing channels, the redundant data, and the un-unique peptides shared by different proteins. Then the data was subject to statistical analysis. For the datasets of non-modified peptides, the quantification was performed at protein level. The log2-values of the measured precursor areas were normalized by the median values across an entire labelling experiment to correct for protein abundance variation. Peptides from same proteins were merged with the R Rollup function (http://www.omics.pnl.gov) allowing for one-hit-wonders and using the mean of the normalized areas for each peptide. Then the mean over the experimental conditions for each protein in each replicate was subtracted, and data from the three replicates were merged. For the datasets of peptides with PTMs, quantification was performed for each peptide in the same way as described above. Detection of differentially regulated proteins and peptides was performed applying a combination of limma and rank tests [60], Resulting p-values were corrected for multiple test [61]. Statistical methods were described in more detail in [62]. Perseus was also used to visualize the statistical results [63]. Very stringent criteria was used to define proteins and modified peptides with significant regulation, which should be observed in at least two replicates, with a p-value LJ 0.05, and showed no less than 1.5 fold change in one condition compared to the other two conditions. We applied fuzzy c-means clustering analysis [64] of peptides with PTMs.

Motif analysis of PTM sites was performed using the Motif-X online software (http://motif-x.med.harvard.edu/motif-x.html) with possibility threshold of P < 10^−6^ and relative occurrence rate threshold of 3% as previously described[65, 66]. Network analysis was performed using Cytoscape (3.4.0 version) and related Apps, including clusterMarker and StringApp with a high confidence setting (0.7), where confident associations were shown with connecting lines. The sub-clusters were generated using Markov CLustering Algorithm (MCL) in clusterMarker app. Ingenuity pathway analysis (IPA) was performed to reveal the protein interaction and signaling pathways by using data source from specific tissue and cell lines (the pancreas, beta cell, and pancreatic cancer cell lines) at high confidence level in order to increase the specificity and decrease the complexity.

## ACKNOWLEDGEMENTS

This work was supported by the postdoctoral fellowship (H.H) from the Danish Diabetes Academy financed by the Novo Nordisk Foundation, the Lundbeck Foundation (M.R.L - Junior Group Leader Fellowship) and by a generous grant from the VILLUM Foundation to the VILLUM Center for Bioanalytical Sciences at the University of Southern Denmark. GP is supported by FAPESP (2014/06863-3), CNPq (441878/2014-8).

## AUTHOR CONTRIBUTIONS

H.H. and M.R.L. conceived and designed all the experiments, H.H. and M.R.L. wrote the paper. H.H. performed the experiments and data analysis. L.D. performed the data analysis. P.S.L. performed cell culture experiment. G.P. provided technical advice and revised the paper.

## Supplementary figures and tables

**Supplementary Figure S1:**
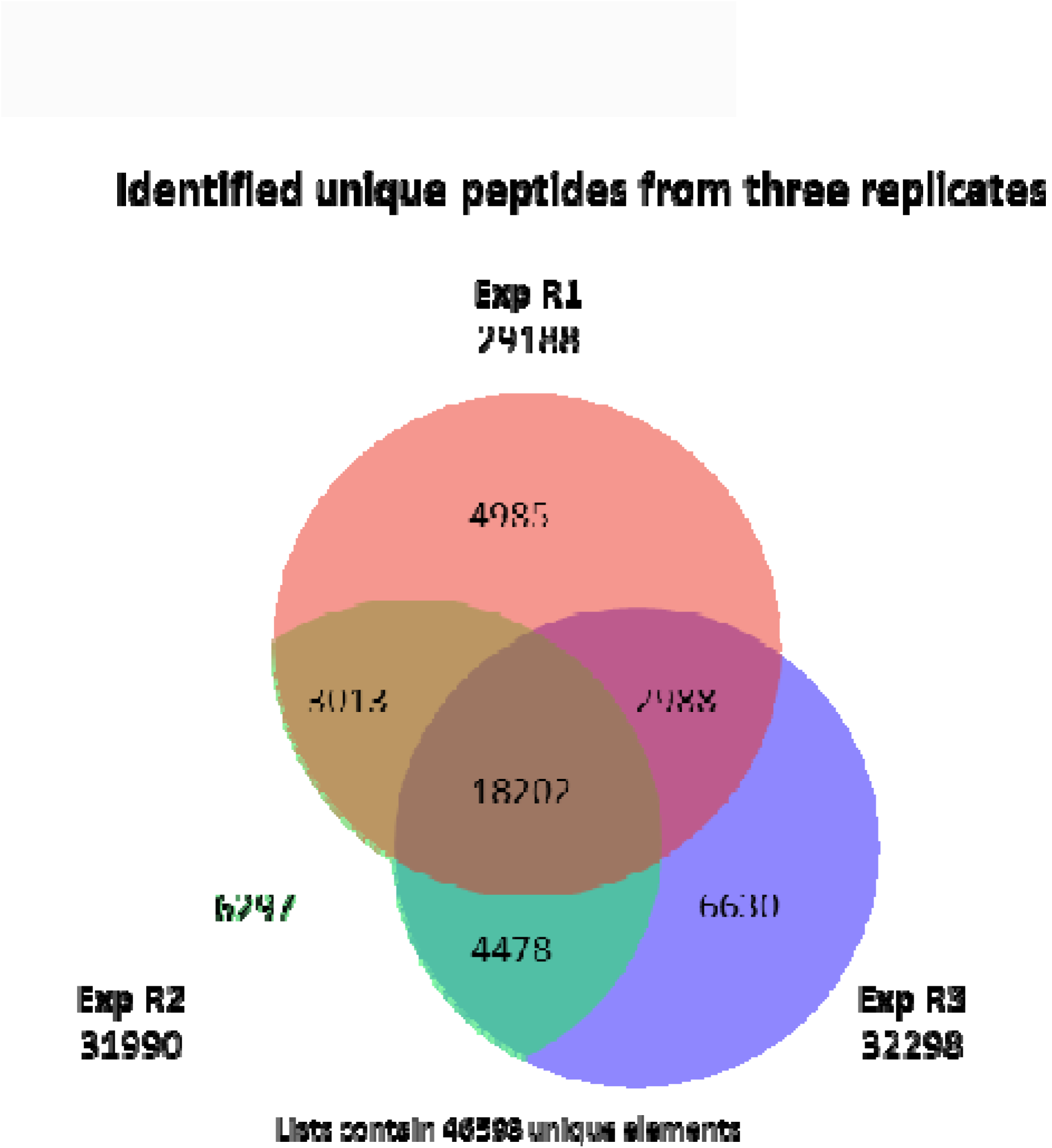
The Venn diagram of identified unique peptides between three experimental replicates from CP fractions.

**Supplementary Figure S2:**
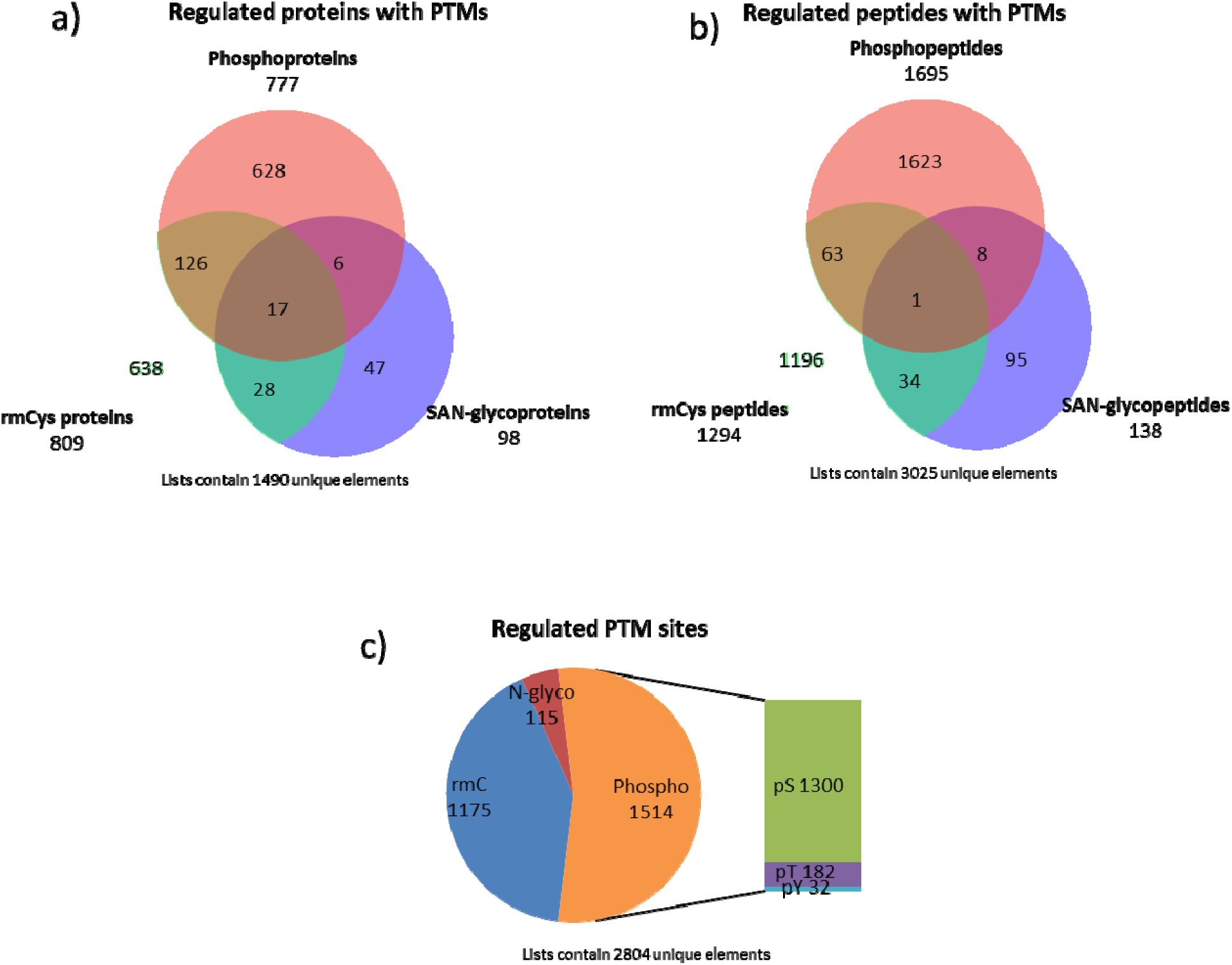
The statistical summary of regulated peptides and proteins with PTMs. (a) The Venn diagram of proteins with regulated rmCys, phosphorylation and SAN-glycosylation. (b) The Venn diagram of regulated rmCys peptides, phosphopeptides and SAN-glycopeptides. (c) The distribution of regulated PTM sites.

**Supplementary Figure S3:**
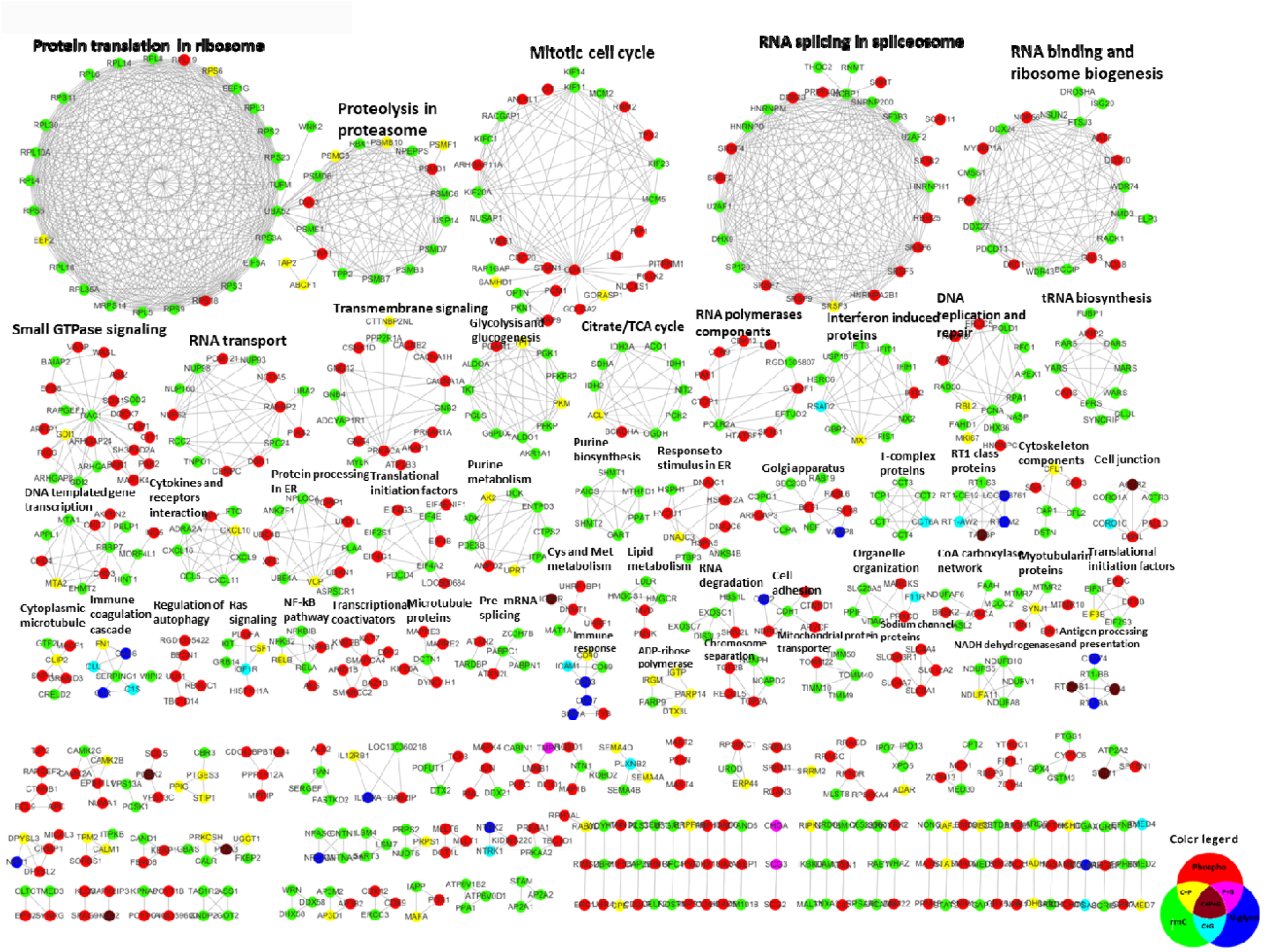
The Markov CLustering Algorithm (MCL) sub-networks of proteins with regulated PTM sites. The MCL sub-clusters were generated based on the STRING network analysis of all proteins with regulated PTM sites using MCL in Cytoscape, and the interesting sub-networks were manually annotated based on their gene ontology or related pathway information. The node color indicated the identified type of regulated PTMs in the protein as presented in color legend.

**Supplementary Figure S4:**
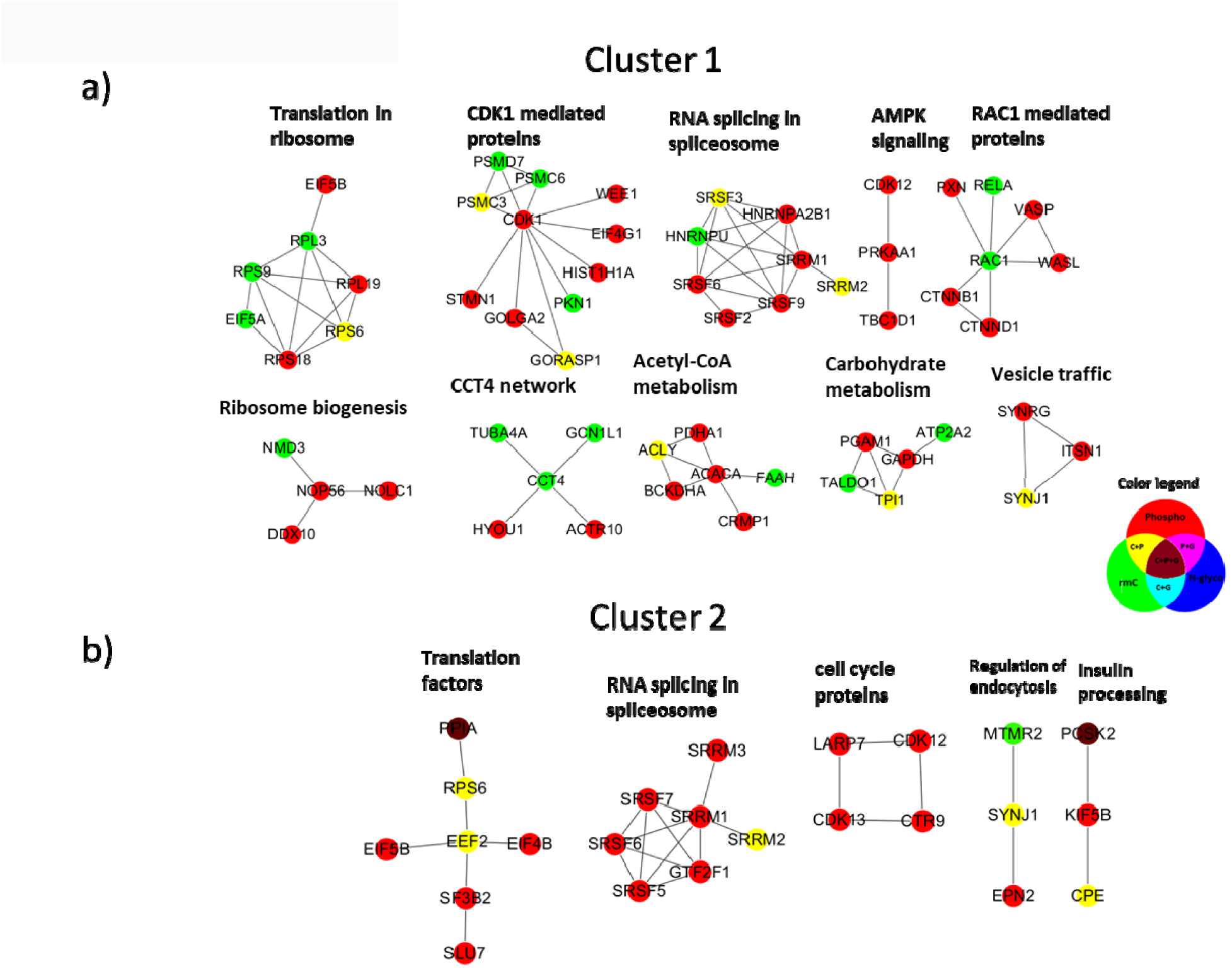
The MCL sub-networks of proteins with regulated PTM sites from fuzzy c-means cluster 1 (a) and 2 (b).

**Supplementary Figure S5:**
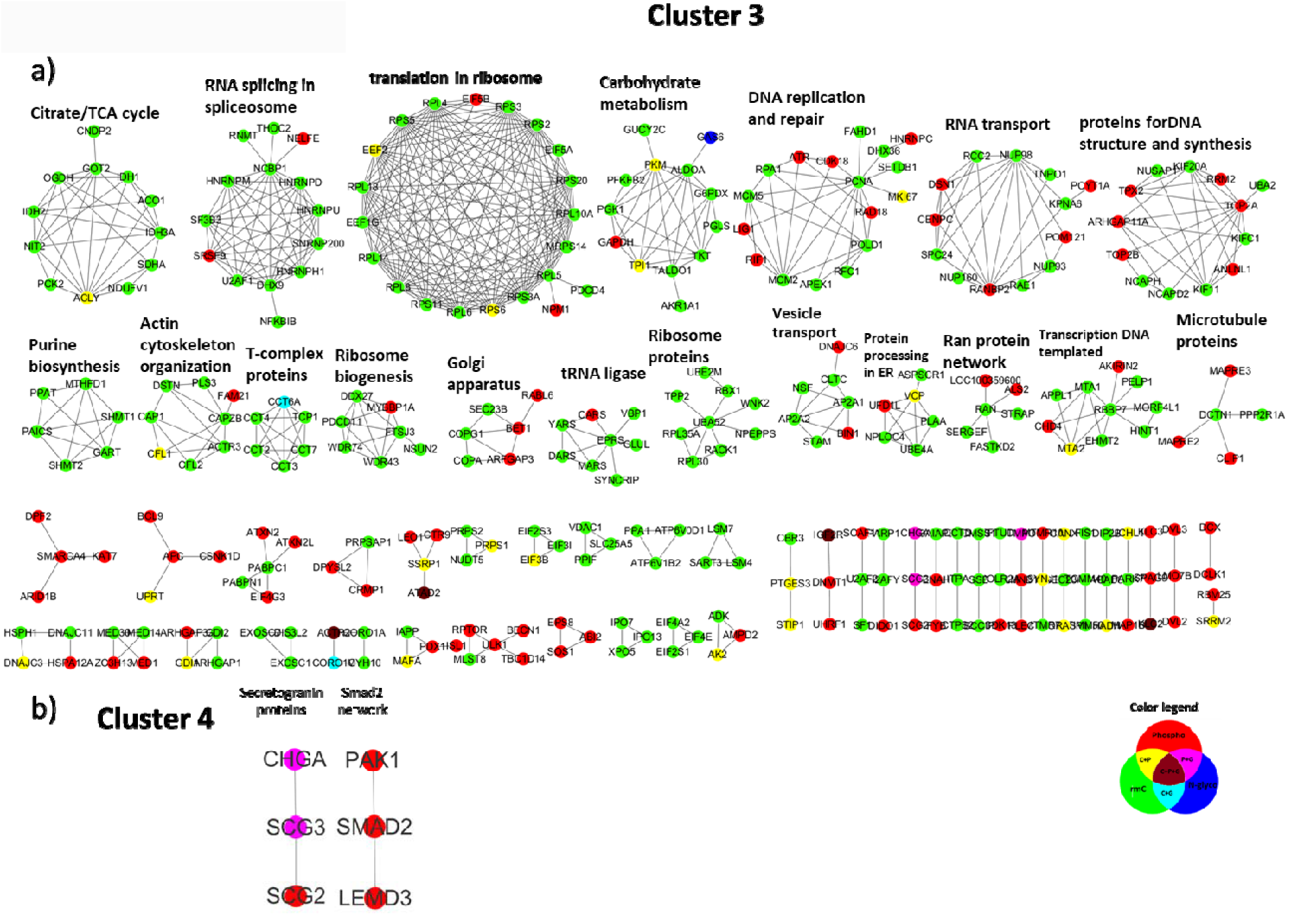
The MCL sub-networks of proteins with regulated PTM sites from fuzzy c-means cluster 3 (a) and 4 (b).

**Supplementary Figure S6:**
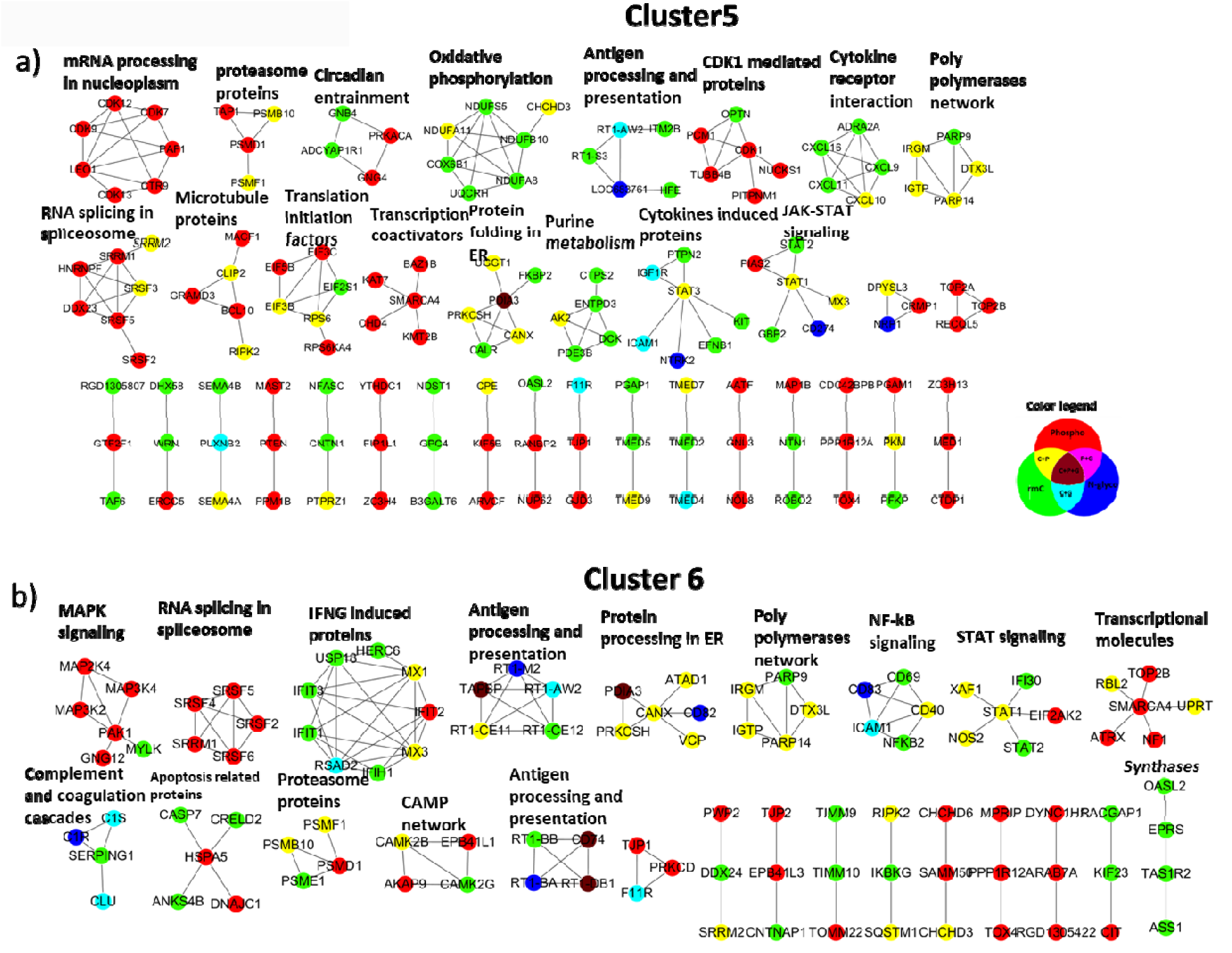
The MCL sub-networks of proteins with regulated PTM sites from fuzzy c-means cluster 5 (a) and 6 (b).

**Supplementary Table S1:** List of identified unique peptides from flow-through of first TiO_2_ enrichment, mainly containing the non-modified peptides.

**Supplementary Table S2:** List of identified unique peptides from the enriched CP and SAN-glyco fractions, mainly containing the peptides with PTMs and some non-modified peptides.

**Supplementary Table S3:** List of identified unique peptides with PTMs from the enriched CP and SAN-glyco fractions.

**Supplementary Table S4:** List of mapped rmCys sites, phosphorylation sites and SAN-glycosylation sites with information of localization on protein, motif window and site annotation retrieved from the Uniprot database.

**Supplementary Table S5:** List of identified motifs for rmCys sites, phosphorylation sites (pS, pT and pY) and SAN-glycosylation sites.

**Supplementary Table S6:** List of significantly regulated proteins at total protein level. The quantification value was the normalized log2 value of intensity.

**Supplementary Table S7:** List of significantly regulated peptides with PTM sites and statistical information. The quantification value was the normalized log2 value of intensity.

**Supplementary Table S8:** List of identified MCL protein sub-clusters from STRING network analysis of all proteins with regulated PTMs. The proteins in each sub-cluster were listed, and only the intense and some interested sub-clusters were annotated.

**Supplementary Table S9:** List of identified 16 most interesting signaling pathways based on the IPA analysis of all identified proteins with regulated PTMs.

## Abbreviations

Cys: Cysteine
CysPAT: Cysteine-specific Phosphonate Adaptable Tag
TiO_2_: titanium dioxide
PTM: post-translational modification
TNF-α: Tumor necrosis factor-α
IFN-γ: interferon-γ
TEAB: triethylammonium bicarbonate
rmCys: reversibly modified Cysteine
TCEP: tris-(2-Carboxyethyl) phosphine
NEM: N-Ethylmaleimide
ACN: acetonitrile
LC: Liquid chromatography
MS: mass spectrometry
HILIC: hydrophilic interaction liquid chromatography
GO: Gene ontology
STAT: signal transducer and activator of transcription proteins
NF-κB: nuclear factor-kappa B
NOS2: inducible NO synthase

